# Environmentally regulated clonal-aggregative multicellularity in a choanoflagellate

**DOI:** 10.1101/2024.03.25.586565

**Authors:** Núria Ros-Rocher, Josean Reyes-Rivera, Uzuki Horo, Chantal Combredet, Yeganeh Foroughijabbari, Ben T. Larson, Maxwell C. Coyle, Erik A.T. Houtepen, Mark J. A. Vermeij, Jacob L. Steenwyk, Thibaut Brunet

## Abstract

Multicellularity evolved multiple times independently during eukaryotic diversification^1–4^. Two distinct mechanisms underpin multicellularity^5^: clonality (serial cell division without sister-cell separation) and aggregation (whereby independent cells assemble into a multicellular entity). Clonal and aggregative multicellularity are traditionally considered mutually exclusive^1,6–9^, with rare exceptions^10^, and evolutionary hypotheses have addressed why multicellularity might diverge toward one or the other extreme^3,4^. Both animals and their sister group, the choanoflagellates, are currently only known to acquire multicellularity clonally^4,11–13^. Here, we show that the choanoflagellate *Choanoeca flexa*^14^ forms motile and contractile cell monolayers (or “sheets”) through multiple mechanisms: *C. flexa* sheets can form purely clonally, purely aggregatively, or by a combination of both processes. We characterise the life history of *C. flexa* in its natural environment – ephemeral splash pools on the island of Curaçao – and show that *C. flexa* undergoes reversible transitions between unicellularity and multicellularity during cycles of evaporation and refilling. Different splash pools house genetically distinct strains of *C. flexa,* between which aggregation is constrained by kin recognition^15–18^. We show that clonal-aggregative multicellularity serves as a versatile strategy for the robust re-establishment of multicellularity in this variable and fast-fluctuating environment. Our findings challenge former generalisations about choanoflagellates and expand the option space of choanozoan multicellularity.

---

Multicellularity has evolved independently more than 45 times across eukaryotes^19^, with multiple independent origins of both clonal and aggregative multicellularity^7,20^. Efforts to reconstruct the origin of animal multicellularity have benefited from the study of their closest living relatives, the choanoflagellates (**Figure 1A**)^21–24^. Choanoflagellates are bacterivorous aquatic microeukaryotes bearing an apical flagellum surrounded by a collar of actin-filled microvilli (**Figure 1B-C**)^25^. Moreover, many choanoflagellate species display facultative multicellularity^25^. The best-characterised choanoflagellate, *Salpingoeca rosetta*, forms colonies exclusively clonally^26^, and clonal multicellularity has classically been assumed to be a general feature of choanoflagellates^4,25,27^. However, this assumption remains to be tested across choanoflagellate diversity. Interestingly, while animal multicellularity is purely clonal, other close relatives of animals besides choanoflagellates exhibit diverse forms of multicellularity, including aggregation in filastereans^28–31^ as well as cellularization of multinucleated cells^32,33^ and cleavage-like serial cell divisions in ichthyosporeans^33–36^.

**Figure 1.**
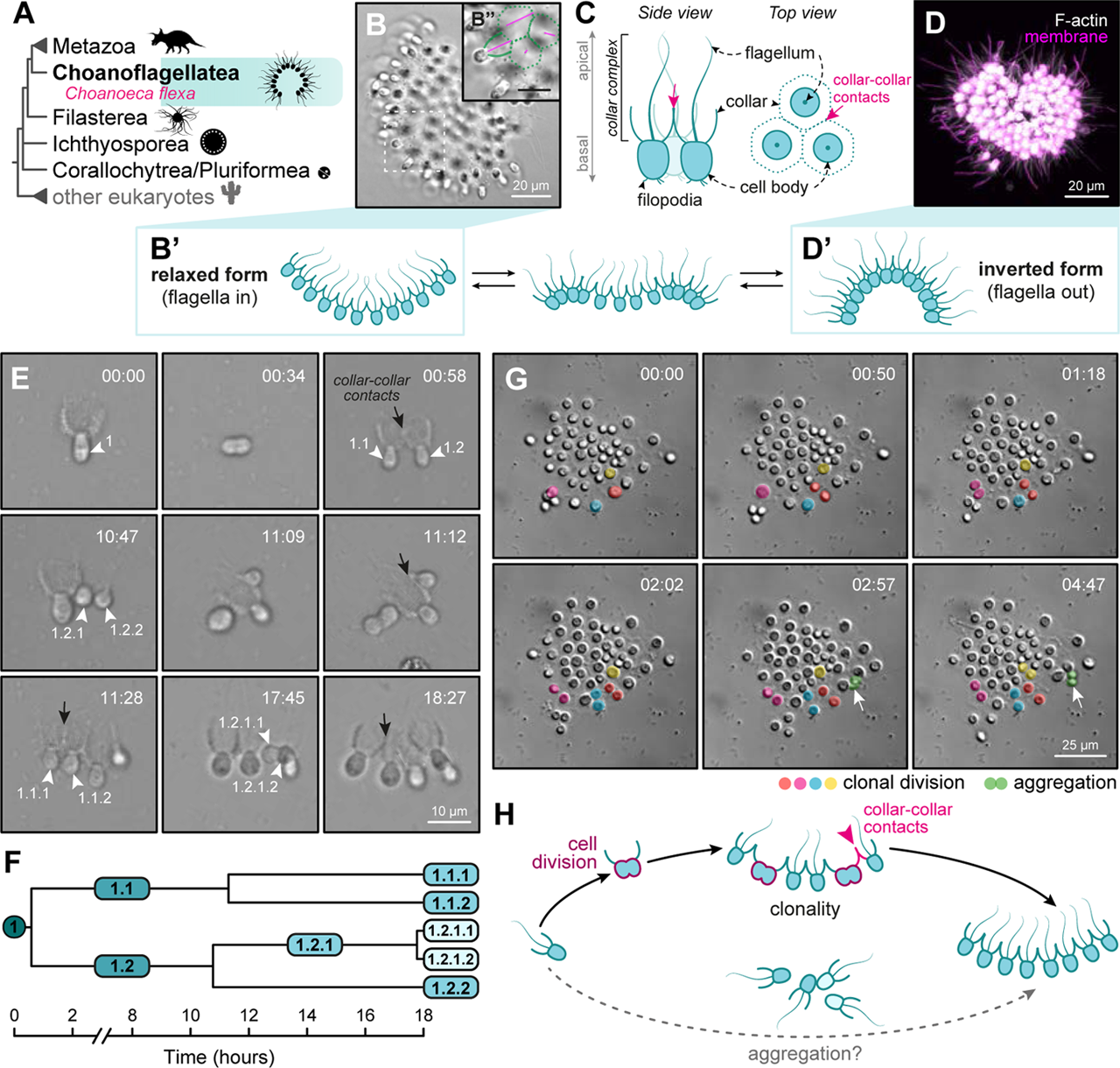
*Choanoeca flexa* sheets can form clonally but display aggregative features. (A) Choanoflagellates (turquoise) are the sister group to animals (Metazoa). The phylogenetic relationships depicted are based on several recent phylogenomic studies^14,97^. Uncertain positions are represented with polytomies. (B) Brightfield image of a *C. flexa* multicellular colony (“sheet”) in its relaxed conformation (B’). B”, dashed square: zoom-in showing flagella (magenta pseudocolour) and direct collar-collar contacts between cells (green pseudocolour). Scale bar in B’’: 10 µm. (C) Diagnostic morphological features of a choanoflagellate cell. *C. flexa* cells within a sheet are linked by their collars (magenta arrow). (D) 3D reconstruction of an Airyscan confocal Z-stack of a fixed sheet exhibiting an inverted conformation (D’), with cell bodies stained with a membrane/cytoplasmic dye (FM 1-43FX, magenta, which distributes to the membrane and cytoplasm of cells following fixation), and collars stained with a filamentous actin (F-actin) dye (Phalloidin-Rhodamine, white). (E) Stills from a brightfield timelapse movie of clonal *C. flexa* sheet formation by serial cell division from a single *C. flexa* swimmer cell (white arrowhead). After each division, the sister cells remain adhered to each other by direct collar-collar contacts between cells (black arrow). Note that cells retract their flagellum during division. Time scale hh:mm. (F) Cell lineage tracing as a function of time in E shows that cells divide asynchronously during colony formation, taking ∼8-10 hours between each round of division. (G) Stills from a brightfield timelapse movie depicting a medium-sized *C. flexa* sheet (flagella-in conformation) that expands in cell number both by cell division (pseudocolours in orange, pink, blue and yellow) and by cellular aggregation (white arrow, pseudocolour in green). Time is hh:mm. (H) Schematics of *C. flexa* clonal multicellularity observed under laboratory culture conditions: unicellular flagellate (swimmer) cells can divide clonally to initiate a colony; in turn, colonies can increase in cell number by clonal division. Sister cells resulting from cell division adhere to each other through direct collar-collar contacts between cells (magenta arrowhead). The hypothesis that *C. flexa* sheets might be able to form purely by aggregation is tested in Figure 2. Figure related to **Figure S1** and **movies S1-S3**.

Here, we describe an unusual mode of multicellularity in the choanoflagellate *Choanoeca flexa*^14^ that challenges prevailing assumptions about choanoflagellates. We show that *C. flexa* colonies can form by serial cell division, aggregation, or a combination of both, a mechanism we refer to as ‘clonal-aggregative multicellularity’. We propose that this mode of multicellularity represents an adaptation to the dynamic natural environment of *C. flexa*: ephemeral splash pools that undergo extreme salinity fluctuations during natural cycles of evaporation and refilling.

## *C. flexa* sheets can form clonally

*C. flexa* was discovered in 2018 in the form of curved monolayers of polarised cells (or ‘sheet colonies’) held together through direct collar-collar adhesions^14^ (**Figure 1A-D**). *C. flexa* sheets can reversibly invert their curvature in response to light-to-dark transitions, switching between a feeding state (relaxed form, flagella-in; **Figure 1B**) and a swimming state (inverted form, flagella-out; **Figure 1D**)^37,38^. In an earlier study, we established stable cultures of *C. flexa* sheets from a single isolated cell, indicating that sheets can arise from individual cells^37^. Nevertheless, the mechanisms that establish multicellularity in *C. flexa* remain unknown.

To understand *C. flexa* colony formation, we isolated and monitored single cells from mechanically disassembled sheets by time-lapse microscopy. In this context, we observed clonal formation of sheets by serial cell division of single cells (**Figure 1E-F; movie S1**): cells divided asynchronously every ∼8-10 hours, and sister cells remained attached to each other by collar-collar contacts. This resulted in a monolayer of polarised cells with the signature curved morphology of *C. flexa* sheets (**Figure 1B,E; movie S1**). These observations show that small *C. flexa* sheets (up to 7 cells) can form clonally. To test whether clonality can contribute to the further growth of *C. flexa* colonies, we monitored small and medium-sized colonies (from 6 to 46 cells) by time-lapse microscopy and observed clonal expansion by cell division both at the core and the periphery of the sheets (**Figures 1G** and **S1; movies S2-S3**). However, and unexpectedly, we also captured instances of free-swimming single cells and doublets meeting colonies, attaching to their periphery, reorienting to align their main apico-basal polarity axis with neighbouring cells, and seemingly integrating into the sheet (**Figures 1G** and **S1; movies S2-S3**). These results suggested that *C. flexa* colonies could form clonally but might also expand by aggregation. This motivated us to test whether *C. flexa* might be capable of purely aggregative multicellularity (**Figure 1H**).

## *C. flexa* sheets can form by aggregation

To test whether *C. flexa* colonies can form by aggregation, we mechanically disassembled colonies into free-swimming single cells and performed live imaging of the dissociated cells (**Figure 2A-D; movies S4-S6**). Surprisingly, *C. flexa* single cells aggregated within minutes into cell doublets connected by collar-collar contacts (**Figure 2C; movie S5**). Doublets grew into larger groups of cells both through the incorporation of additional solitary cells and through fusion between groups (**Figure 2D; movie S6**). Early aggregates were often irregular in shape, reflecting initial collision and adhesion in diverse, random orientations. Aggregates later underwent morphological maturation through cellular rearrangements and reorientation into polarised monolayers with canonical *C. flexa* sheet morphology (**Figure 2D; movie S6**).

**Figure 2.**
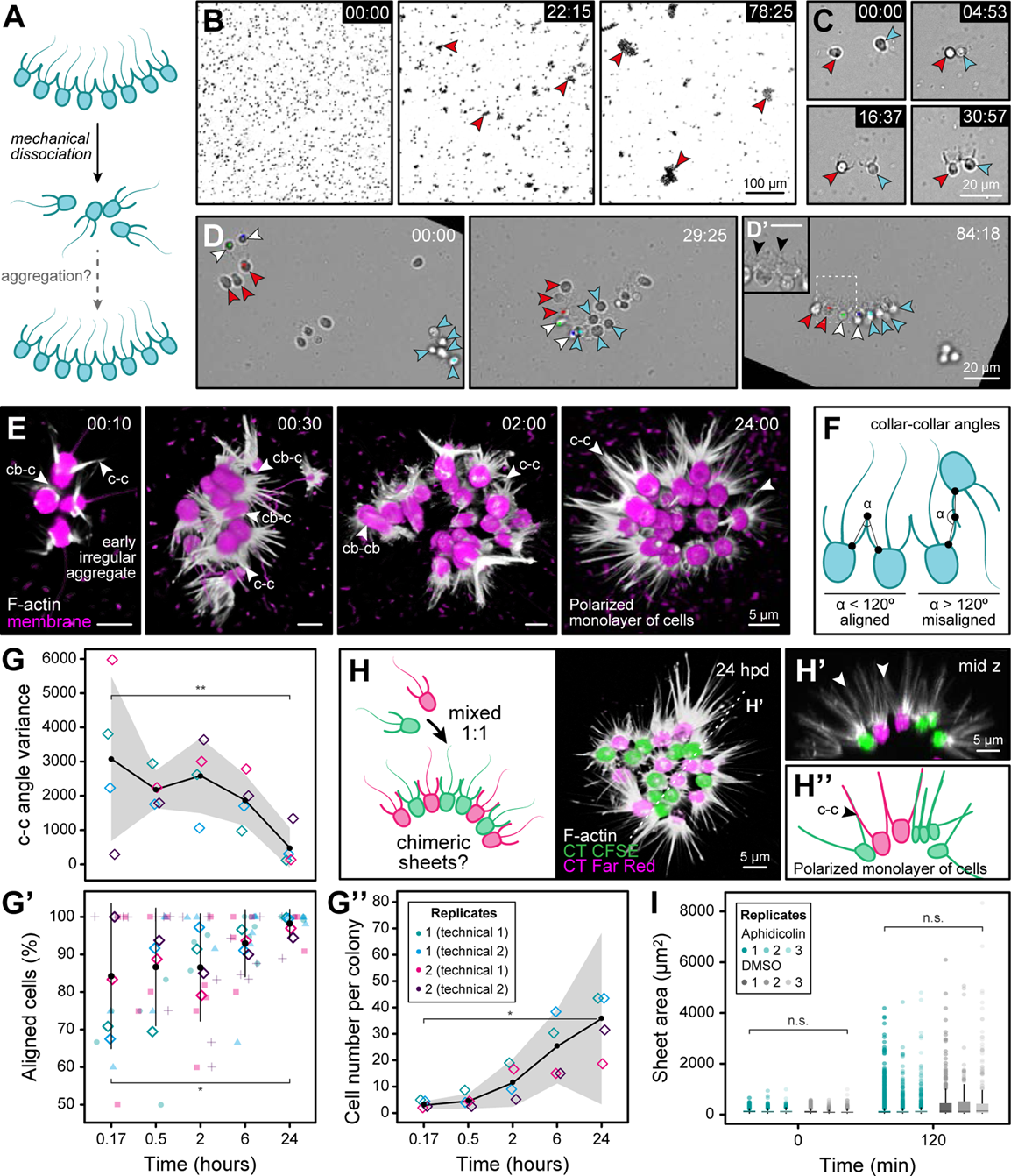
*C. flexa* sheets can form by aggregation. (A) Experimental pipeline. Mechanical dissociation of colonies into single cells is followed by live imaging. (B) Dissociated flagellate cells form aggregates (red arrowheads) within minutes. Stills from Movie S4. Upper right: time. (C) Two dissociated flagellate cells (red and blue arrowheads) aggregating into a doublet connected by collar-collar contact. Stills from Movie S5. Upper right: time. (D) A cell doublet (white arrowheads) and two small sheets (red and blue arrowheads) at t=0 (left, pre-aggregation) aggregate into a larger sheet. The aggregate initially comprises cells with diverse orientations (middle, irregular aggregate) but later matures into a polarised monolayer showing cells with the same apico-basal orientation connected by collar-collar contacts (right, mature aggregate). Stills from Movie S6. Upper right: time. (D’) Dashed square: closeup of the area in the white dotted square in D, showing collar-collar contacts (black arrowheads). Scale bar: 10 µm. Timescales in B-D: mm:ss. (E) 3D reconstruction of Airyscan confocal z-stacks of dissociated flagellate cells fixed after 10 minutes (n=11), 30 minutes (n=14), 2 hours (n=17), and 24 hours (n=13) of aggregation, stained for membrane/cytoplasm (FM 4-64FX, magenta) and F-actin (phalloidin, white). Note that cells frequently show unaligned apico-basal polarity and diverse cell orientations at early timepoints, but almost exclusively collar-collar contacts at late timepoints (white arrowheads; cb-c: cell body-collar contacts, cb-cb: cell body-cell body contacts, c-c: collar-collar contacts). Time scale: hh:mm. (F) Schematics of the morphological metrics quantified from the cells in E: adhesion angle between collar-collar contacts of neighbouring cells (α) and proportion of cells with aligned apico-basal polarity. (G) Quantification of collar-collar angle variance between cells during aggregation in E. (G’) Quantification of percentage of aligned cells within colonies in E. (G’’) Quantification of cell number per colony during aggregation in E. Black circles: mean; error bars and grey ribbon: standard deviation; diamonds: mean values of each independent replicate. (H) (Left panel) Dissociated single flagellate cells labelled with CellTrace CFSE (green) or CellTrace Far Red (magenta) and mixed at a 1:1 ratio will form chimeric sheets 24 hours post-dissociation (hpd) if aggregation occurs. (Right panel) 3D reconstruction of Airyscan confocal z-stacks of a sheet formed by aggregation from dissociated flagellated cells labelled as depicted before and fixed 24 hpd, with additional F-actin staining (phalloidin, white). (H’) Mid z cross-section (dashed line) in H. Note that cells within the chimeric sheet are connected by collar-collar contacts (white arrowheads) and maintain their apico-basal polarity. (H’’) Schematics of H’, showing a polarised monolayer of cells and collar-collar contacts (c-c) of the chimeric sheet in H. (I) Quantification of particle area during an aggregation time course of dissociated single cells pre-treated with 17 µg/mL Aphidicolin overnight or the equivalent volume of a DMSO (dimethyl sulfoxide) control (n=9). Statistics in G-G’’ (comparing t=10 min and t=24 h) and I (comparing average area of Aphidicolin-treated versus DMSO control at each timepoint) by the Mann-Whitney U test (* for p<0.05; ** for p<0.01; *** for p<0.001; n.s., non-significant). Statistics in G’ by linear regression: adjusted R^2^ = 0.081, p-value= 0.042. Figure related to **Figures S2-S5** and **movies S4-S13**.

To quantify the dynamics of aggregation and maturation, we fixed sheets at successive stages of aggregation, followed by membrane and F-actin staining and Airyscan confocal imaging (**Figures 2E-G** and **S2; movies S7-S11**). We quantified two morphological metrics: the adhesion angle between the collars of neighbouring cells (or collar-collar angle) and the proportion of cells with aligned apico-basal polarity (**Figures 2F** and **S2A**). Both metrics initially showed high variance, consistent with variable cell orientations at early stages, and progressively converged towards stereotypical values during maturation (**Figures 2E-G** and **S2; movies S7-S11**). Moreover, early aggregates showed diverse types of intercellular contacts, including collar-collar, collar-cell body, and cell body-cell body contacts (**Figures 2E** and **S2B**). By contrast, mature colonies resulting from 24 hours of aggregation were polarised monolayers of cells connected almost exclusively by collar-collar contacts (**Figures 2E** and **S2B; movie S11**). At that stage, colonies comprised about 50 cells on average and up to 120 cells (**Figures 2G’’** and **S2D**). This size further confirmed that they could not have formed exclusively through cell division within 24 hours, given the cell cycle duration in *C. flexa* (at least 8 hours; **Figure 1F**), which would allow for sheets of at most 16 cells, assuming maximal and synchronised proliferation.

To independently confirm that the regular sheets observed 24 hours post-dissociation (hpd) resulted from the maturation of early aggregates, we performed aggregation assays by mixing two populations of dissociated single cells stained with different fluorophores (**Figures 2H** and **S3; movies S12-S13**). Cells of both colours assembled within minutes into irregular chimeric aggregates (**Figure S3A; movie S12**). After 24 hours, we observed chimeric polarised monolayers composed of cells of both colours (**Figures 2H** and **S3B-C; movie S13**), confirming their aggregative origin. Furthermore, treatment with the cell cycle inhibitor aphidicolin did not abolish aggregation (**Figure 2I** and **S4**), confirming that aggregation can occur independently of cell division. Finally, we sought to determine whether aggregation was an active process or whether it might simply result, at least at early stages, from passive cell stickiness. While live cells readily aggregated, fixed cells did not (even under orbital agitation forcing cell encounters), suggesting that aggregation requires live cells and is thus an active process (**Figure S5**).

Taken together, these data indicate that *C. flexa* sheets can form by aggregation. This process is independent of cell division, is active, and occurs in multiple steps: random cell collisions initially form irregular clumps of variable morphology, which then mature into regular, polarised monolayers of cells via cell reorientation.

The discovery of clonal-aggregative multicellularity in *C. flexa* contrasts with prevailing assumptions about choanoflagellates, which – based largely on studies of the model species *S. rosetta*^39^ – were thought to acquire multicellularity only clonally. Notably, clonal and aggregative multicellularity usually occupy different ecological niches: aggregation is often an ‘emergency response’ to sudden stress^3,40^, whereas clonality requires sufficiently stable environmental conditions to sustain cell division. This prompted us to investigate the natural environmental context in which *C. flexa* might employ clonal-aggregative multicellularity.

## Occurrence of multicellular *C. flexa* in the field is limited by salinity

*C. flexa* was originally discovered in its multicellular form on the tropical island of Curaçao^14^ (**Figure 3A**). Unlike other multicellular choanoflagellates^22,41^, *C. flexa* has been repeatedly re-isolated in the field since its discovery in 2018 and can thus be studied in its native biotope. *C. flexa* sheets are found on the windward, northern part of the island, in splash pools that undergo natural cycles of evaporation and refilling^42,43^ (**Figures 3A-C** and **S6A-B; movie S14**). Water-filled splash pools gradually evaporate, leading to increasing salinity and, occasionally, complete desiccation (**Figure 3C**). Splash pools are subsequently refilled by waves, splash, or rain, restoring lower salinity levels (**Figures 3C** and **S6B; movie S14**). As a consequence, splash pools are ephemeral habitats in which organisms are exposed to extreme and recurrent hypersaline and hyperosmotic stress^43–46^. We therefore set out to investigate how this dynamic habitat might influence the life history and multicellularity of *C. flexa*.

**Figure 3.**
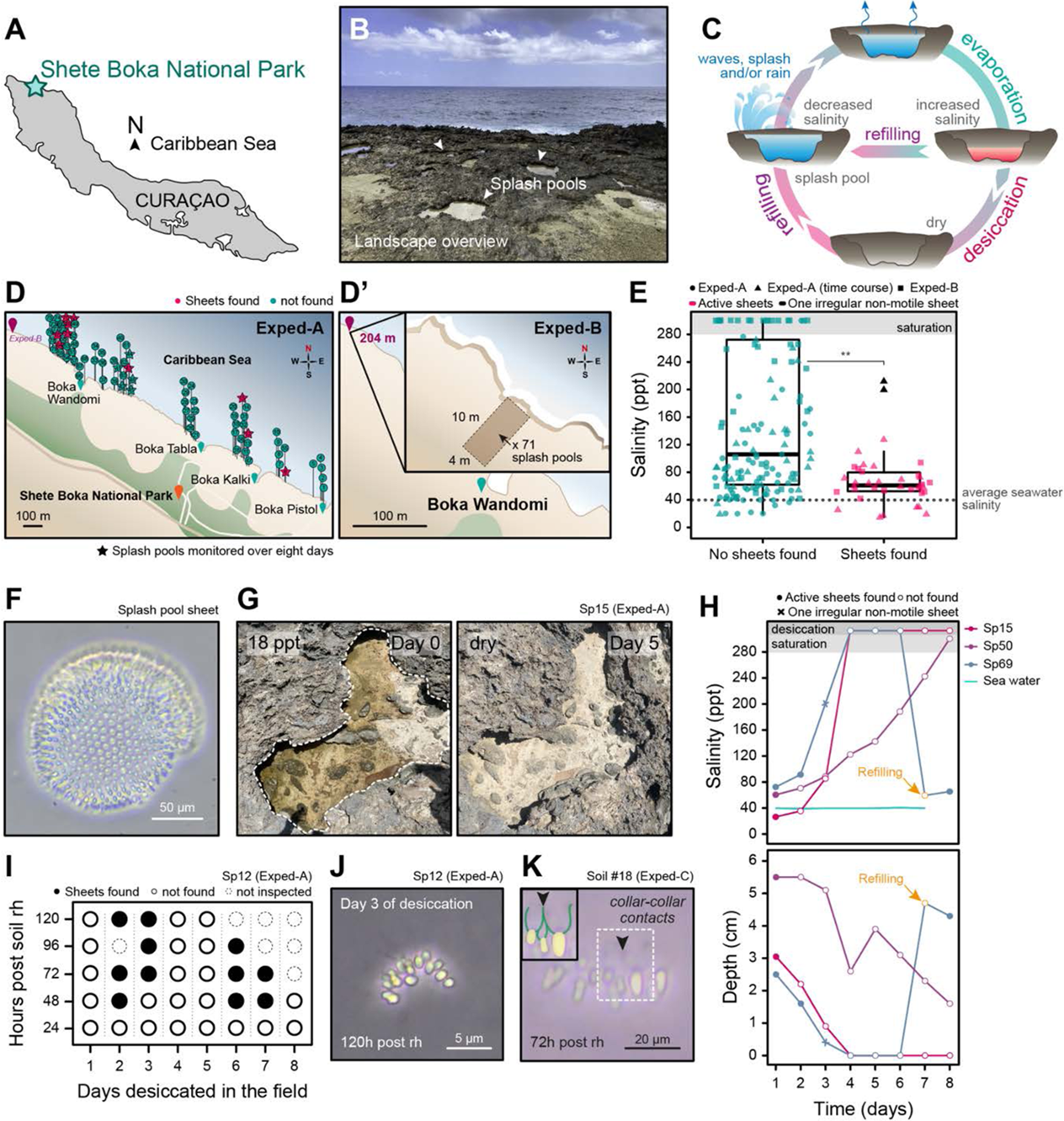
Natural evaporation-refilling cycles constrain the occurrence of multicellular *C. flexa* in the wild. (A) Map of Curaçao and location of Shete Boka National Park (turquoise star) where fieldwork data were collected in Exped-A and Exped-B (12° 22’ 5.718 “N, 69° 06’ 56.916” W). (B) Representative photograph of the landscape in Shete Boka, including splash pools where *C. flexa* sheets can be found (white arrowheads). (C) Schematic of the natural cycle of splash pool evaporation, desiccation, and refilling. (D) Maps showing the locations of sampled splash pools in Exped-A and in Exped-B. In Exped-A (D), samples were collected from splash pools along ∼2 km of Shete Boka coastline (n=79). 10 splash pools with sheets and 5 without sheets on Day 1 were randomly selected for daily monitoring during eight days (stars). Colours indicate whether sheets were found (magenta) or not found (turquoise) on Day 1. In Exped-B (D’), a random number generator was used to select the randomised sampling location (purple pin, 204 m upstream of Boka Wandomi). Samples were collected from splash pools within a 10 m x 4 m area (n=71). (E) Distribution of salinity of splash pools surveyed in Exped-A (circles), Exped-A time course (triangles), and Exped-B (squares). For Exped-A time course, all measurements are shown, except for those where splash pools were dry. The observed natural limit of salinity where active sheets were found (128 ppt) and the average seawater salinity measured in the *bokas* are indicated with grey dotted lines. Grey area: salinity saturation (>280 ppt). (F) Brightfield image of a sheet observed in a splash pool sample (n=18). (G) Representative images of a splash pool near Boka Kalki (Sp15) followed for eight days, showing recorded salinity in the upper left. This splash pool was partially evaporated on day 0 (left) and completely desiccated on day 5 (right). Dashed line: splash pool outline. (H) Salinity (upper panel) and depth (lower panel) measurements in three representative splash pools followed over eight days (n=15). Shown are one splash pool that experienced evaporation but not complete desiccation (Sp50), one splash pool that experienced complete desiccation (Sp15), and one splash pool that experienced both desiccation and refilling (Sp69). (I) Recovery of sheets from soil samples collected from a splash pool (Sp12 from Exped-A) when desiccated. Soil samples were collected every day for eight days and were independently rehydrated in the laboratory with filtered natural seawater. Each rehydrated soil sample was monitored over five days for sheet re-appearance. rh: rehydration. Rehydration experiments were performed from soil collected in n=6 independent splash pools, with a minimum of three independent rehydrations each. (J) Brightfield image of a soil-recovered sheet (same experiment as in I) collected after 3 days of desiccation and rehydrated in the laboratory. (K) Brightfield image of a soil-recovered sheet independently collected and rehydrated from a splash pool sample (Soil #18) in Exped-C, showing collar-collar contacts (black arrowhead). Dashed square: detail of cell-cell contacts by the collars (black arrowhead). Green pseudocolour: collars; yellow pseudocolour: cell bodies. Rehydration experiment performed from soils collected in n=26 independent splash pools. Time post-rehydration in J-K is depicted in the lower left. Figure related to **Figures S6-S9**, **movies S14-S16**, **Tables S1-S2** and **Supplementary Files S1-S4**.

We focused our studies on Shete Boka National Park, where *C. flexa* is consistently found (**Figure 3A**). We measured salinity and scored the presence of *C. flexa* sheets in 150 splash pools in two different field expeditions, Exped-A and Exped-B (**Figures 3D** and **S6; Table S1**). During Exped-A, we sampled 79 splash pools (numbered Sp1 to Sp79) along approximately 2 kilometres of coastline (**Figure 3D**; see Materials and Methods for details). While seawater collected from adjacent inlets (or *bokas*) had a stable salinity of ∼40 parts per thousand (ppt; **Table S1**), splash pool salinity ranged from below that of average seawater salinity (15 ppt) to saturation (≥ 280 ppt) (**Figures 3E** and **S6D; Table S1**). In 10 of the 79 splash pools, we found choanoflagellate sheets that we identified as *C. flexa* based on morphology, inversion behaviour, and 18S ribosomal DNA (rDNA) sequencing (**Figures 3F** and **S7; movie S15; Tables S1-S2; Supplementary files S1-S4**). *C. flexa* sheets were not observed in the other 69 splash pools. Interestingly, although the salinity of surveyed splash pools ranged from 15 to 280 ppt, *C. flexa* sheets were predominantly found in splash pools with ∼2-fold seawater salinity during Exped-A (**Figures 3E** and **S6D; Table S1**). To independently test whether the presence of multicellular *C. flexa* was constrained by an upper salinity limit, we exhaustively sampled all splash pools within a 4-meter by 10-meter quadrant during a second expedition (Exped-B; n=71 splash pools, numbered M1 to M71; **Figure 3D’** and **S6C; Table S1**). We observed sheets in 14 splash pools and not in 57 others, with a maximum salinity threshold of 94 ppt for sheet occurrence (**Figures 3E** and **S6D**). Across both expeditions, *C. flexa* sheets were found in splash pools with an average salinity of 62.1±24.8 ppt (equivalent to 1.55-fold seawater salinity; **Figures 3E; Table S1**), which was significantly lower than the salinity of splash pools in which sheets were not observed (146.3±95.7 ppt; p=1.7e-06 by the Mann-Whitney U test; **Figures 3E; Table S1**). Across both expeditions, we never observed actively swimming and inverting *C. flexa* sheets in splash pools exceeding 128 ppt (∼3-fold seawater salinity; **Figures 3E** and **S6D; Table S1**).

## Natural evaporation-refilling cycles cause reversible transitions between unicellular and multicellular states in splash pools

We next examined how the natural evaporation-refilling cycles of splash pools impacts the occurrence of sheets. We monitored ten splash pools where *C. flexa* sheets had already been found during Exped-A, along with five additional randomly selected splash pools, once per day during eight days (**Figures 3E,G-H** and **S8**). We recorded salinity and maximum depth, and noted the presence of sheets (**Figures 3E,G-H** and **S8**). We observed a progressive increase in salinity and decrease in depth in all 15 splash pools (presumably due to evaporation; **Figure S8A-B**), with six desiccation events (**Figure S8C-D**) and four refilling events (**Figure S8E-F**; three examples are shown in **Figure 3H**). In all cases, actively swimming and inverting *C. flexa* sheets were no longer observed after salinity exceeded a 128 ppt threshold during gradual evaporation (**Figures 3H** and **S8**), consistent with results from Exped-A and Exped-B (**Figures 3E-H** and **S8**). Interestingly, we observed two isolated colonies with an apparently stressed phenotype (characterized by irregular outlines, loose cell packing, lack of flagellar beating, and absence of inversion behaviour) at 200 and 212 ppt in two different splash pools (Sp64 and Sp69, respectively on Days 4 and 3; **Figures 3H** and **S8E-F; Table S1**). These irregular and inactive colonies were not observed at later time points, suggesting they had either died or dissociated (**Figures 3H** and **S8F; Table S1**). Similarly, gradual evaporation in the laboratory of a natural splash pool sample containing sheets led to sheet disappearance (**Figure S9**). These observations confirmed that the multicellular form of *C. flexa* does not tolerate high salinity.

Notably, in two dry splash pools that underwent refilling during our study (where salinity was restored to ∼50 ppt), sheets were observed 48 hours after refilling (Sp69 and Sp70; **Figures 3H** and **S8E-F**). The newly observed sheets may have originated from the ocean or from neighbouring splash pools. Alternatively, *C. flexa* might have persisted in the soil of desiccated splash pools in a cryptic resistant form, such as unicellular cysts (as described in other choanoflagellates^47–49^). To test this possibility *in situ*, we collected soil samples from six desiccated splash pools in which *C. flexa* had been previously observed, rehydrated them in the laboratory, and monitored the rehydrated samples for several days (**Figure 3I-K; movie S16** and **Table S1**). Sheets were consistently recovered from soil samples from two splash pools after 2-7 days of desiccation in the field followed by 2-3 days of rehydration in the laboratory, suggesting that a resistant form of *C. flexa* can survive complete desiccation (**Figure 3I-K** and **Table S1**).

Given that salinity fluctuations in splash pools seemed to impact *C. flexa* multicellularity in nature, we next set out to investigate the phenotypic response of *C. flexa* to comparable evaporation-refilling cycles in a laboratory context.

## Evaporation-refilling cycles cause reversible transitions between multicellular and unicellular states in the laboratory

To mimic natural evaporation-refilling cycles in the laboratory, we subjected *C. flexa* cultures to gradual evaporation in an incubator (**Figure 4A**), replicating the temperature (30°C) and evaporation rate of natural splash pools (**Figure 4B** and **S10A;** see Materials and Methods). Starting from the standard salinity of artificial seawater (35 ppt; hereafter ‘1X salinity’), this resulted in complete desiccation after four days. Under these conditions, *C. flexa* sheets gradually dissociated into non-motile single cells (**Figures 4C-D** and **S10B; movie S17**), with more than 50% solitary cells observed once salinity crossed the natural limit of active sheet occurrence (128 ppt; **Figures 3E** and **4C-D**). By the time salinity reached saturation, nearly all sheets had dissociated into single cells (**Figures 4C-D** and **S10B; movie S17**). By contrast, control cultures maintained under constant conditions without evaporation remained multicellular throughout the experiment (**Figure S10C**). Finally, to directly test whether the loss of multicellularity was caused by increased salinity, we added seawater salt to *C. flexa* cultures (without evaporation) and similarly observed dissociation of multicellular sheets into single cells (**Figure S11A-C**).

**Figure 4.**
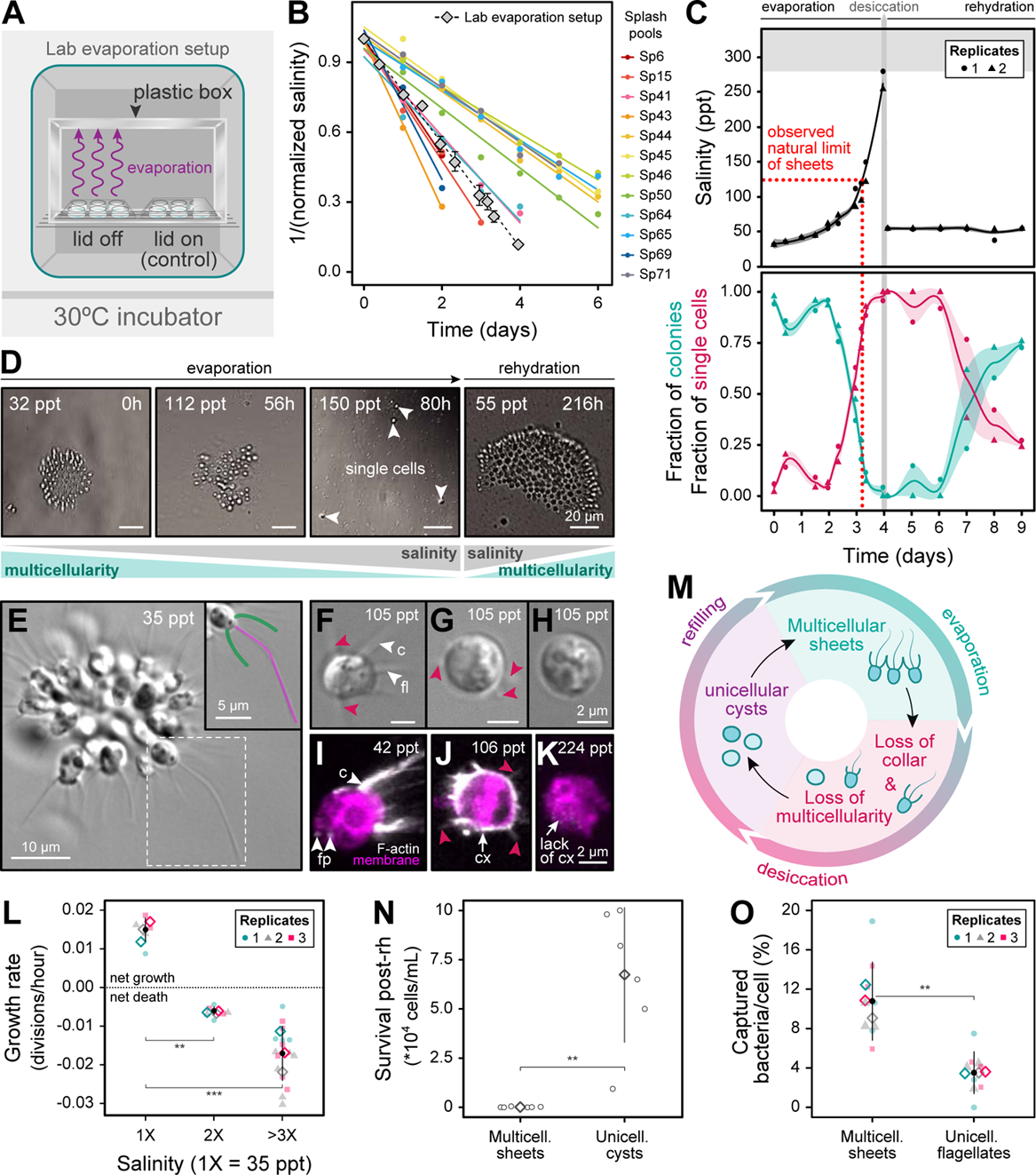
Experimental evaporation-refilling cycles cause reversible transitions in and out of multicellularity. (A) Laboratory experimental evaporation setup in a 30°C incubator. Lid-off plates (gradual evaporation condition) and lid-on plates (no-evaporation control) were placed on a grid to allow air renewal and covered with a plastic box to avoid contamination. (B) Salinity increases over time in 12 gradually evaporating splash pools (data from Exped-A time course; Figures 3H and **S8**) and in the laboratory evaporation setup in A. Splash pools with fewer than three points below saturation are not depicted (Sp12, Sp42, and Sp70; see **Figure S10**). (C) Quantification of salinity and fraction of cells in multicellular versus unicellular forms during a nine-day gradual evaporation time course. Lines: mean values. Ribbon: standard deviation. (D) Micrographs of *C. flexa* sheets during the gradual evaporation experiment in C. Upper left: salinity. Upper right: time. Sheets had completely dissociated into single cells (white arrowheads) after 80 hours. Multicellular sheets were observed again after complete desiccation and rehydration with artificial seawater. (E-H) Brightfield images of *C. flexa* showing morphological changes during gradual evaporation. Upper right: salinity (in ppt). At 1X salinity (35 ppt, E), *C. flexa* occurs in multicellular sheets of flagellates (dashed square). Green pseudocolour: collar; magenta pseudocolour: flagellum. During gradual evaporation (F-H), sheets dissociate into unicellular cysts. Cysts often lack a collar (c) and a flagellum (fl) but can exhibit filopodia-like protrusions (magenta arrowheads). (I-K) Airyscan micrographs of *C. flexa* cells fixed during gradual evaporation stained with a membrane (FM 4-64FX, magenta) and F-actin (phalloidin) dyes, showing salinity (in ppt, upper right). A flagellate cell (I, low-evaporation control) exhibits distribution of F-actin in the collar (c, white arrowhead) and filopodia (fp, white arrowhead). Gradual evaporation triggers a morphological change from a flagellate to a cyst, showing filopodia-like protrusions (magenta arrowheads) and a transient actin cortex (cx) at early stages of evaporation (J, white arrow) which disappears as salinity approaches saturation (K). (L) Growth rate of cells at different salinities during gradual evaporation. Black circles: mean; error bars: standard deviation; diamonds: mean values of independent biological replicates. (M) Schematic summarizing loss of multicellularity and phenotypic changes experienced by *C. flexa* cells during gradual evaporation. (N) Quantification of cell survival after 12 hours of desiccation in multicellular sheets (n=6) to unicellular cysts (n=6). Diamonds: means; error bars: standard deviations.(O) *C. flexa* multicellular sheets are more efficient at capturing bacteria than unicellular flagellates. Number of labelled bacteria captured per flagellate cells in multicellular colonies (n=9) or in cultures of unicellular flagellates (n=9). Black circles: means; error bars: standard deviations; diamonds: means of independent biological replicates. Statistics in L, N, and O by the Mann-Whitney U test (* for p<0.05; ** for p<0.01; *** for p<0.001; n.s., non-significant). Figure related to **Figures S10-S16** and **movies S17-S18**.

We then tested whether single cells resulting from sheet dissociation were viable and capable of surviving desiccation. We rehydrated samples by adding artificial seawater three hours after desiccation, thereby mimicking refilling by waves. This restored salinity to ∼50 ppt and was followed by the reappearance of multicellular sheets within 24 hours post-rehydration, as well as by a continuous increase in the number of colonies over the following days (**Figure 4C-D**). We assessed the mechanism of colony re-formation by time-lapse microscopy of desiccated cultures after rehydration and captured instances of unicellular flagellates engaging in both clonal divisions and aggregation (**Figure S10D; movie S18**). Taken together, our results show that *C. flexa* sheets dissociate into non-motile single cells during gradual evaporation, that these solitary cells can survive complete desiccation, and that sheets reform after rehydration through both clonal division and aggregation. These findings are consistent with our field observations, where we recovered sheets after rehydrating soil samples from dry splash pools (**Figure 3I-K** and **Table S1**), and further support the existence of a desiccation-resistant form of *C. flexa*.

## *C. flexa* sheets undergo dissociation-encystation at high salinity

We next investigated the phenotype of desiccation-induced single cells. In diverse protists, resistance to desiccation is achieved through differentiation into cysts^50–52^ – dormant cells characterized by reduced metabolic activity and proliferation. Encystation often entails significant morphological changes, including cell rounding, flagellar loss, and the formation of a protective cell wall^47,50^. We monitored cellular morphology by DIC microscopy during gradual evaporation and observed asynchronous structural changes that produced a heterogeneous cell population (**Figure 4E-H**). At 3X salinity, most cells had dissociated from their colonies and displayed a rounded cell body lacking microvilli, often missing a flagellum and occasionally bearing filopodia-like protrusions (**Figure 4F-H**). Membrane and F-actin staining of desiccation-resistant cells (hereafter ‘cysts’) confirmed the absence of a collar complex and revealed a transient F-actin cortex, detectable at 3X salinity but lost above 6X (near saturation; **Figures 4I-K** and **S12**). This cortex may help protect the plasma membrane against osmotic stress during early differentiation^53–56^. Cysts did not proliferate: growth was arrested above 2X salinity, with a net cell loss above 3X salinity (**Figures 4L** and **S13**), similar to cell cycle arrest during encystation in other protists^50,57^. Finally, morphometric analysis of cell body and nucleus volume showed that cysts had a larger nucleus-to-cytoplasm ratio compared with flagellates at 1X salinity (**Figure S14**). An increase in the nucleus-to-cytoplasm ratio is frequently associated with cell quiescence^58^, consistent with growth arrest in *C. flexa* cysts. To confirm that these changes were induced by hypersaline stress, we directly increased the salinity of *C. flexa* cultures by adding seawater salts and observed similar morphological changes and arrest in cell growth (**Figure S11**). Thus, the cellular changes induced by hypersalinity in *C. flexa* closely resemble encystation in diverse other protists^50,59^.

Sheet dissociation during encystation is likely caused by retraction of the microvilli that connect neighbouring cells within colonies^14^. We tested this hypothesis by inducing microvillar retraction (independently of encystation) by treatment with the actin-depolymerizing compound latrunculin B, and observed dissociation of colonies within minutes (**Figure S15A-D**). Similarly, latrunculin B treatment of single cells prevented aggregation (**Figure S15E-H**). These results confirm that microvillar integrity is necessary for multicellularity in *C. flexa* and further support the idea that sheets undergo a coupled dissociation-encystation process under hypersaline conditions.

During the natural evaporation-refilling cycles of splash pools, *C. flexa* therefore alternates between multicellular, flagellated sheets (at low salinity) and unicellular cysts (at high salinity), suggesting these two phenotypes each confer an adaptive advantage in their respective environments (**Figure 4M**). We then sought to directly test these putative advantages under laboratory conditions.

## Multicellular sheets and unicellular cysts are respectively advantaged at low and high salinity

If *C. flexa* cysts represent a desiccation-resistant form, we could expect these cells to be more resistant to hypersaline stress and desiccation than the sheets. To test this, we induced the differentiation of *C. flexa* sheets into cysts by gradual evaporation, following the protocol detailed in the previous section (from 1X salinity to complete desiccation over 72 hours). In parallel, we subjected sheets to rapid evaporation, causing desiccation over a period of 20 hours from 1X salinity (see Materials and Methods). Under these conditions, cells did not acquire a cyst-like morphology but instead retained their flagellum, collar, and multicellular morphology, even when completely desiccated (**Figure S16A**). This observation suggests that the formation of cysts requires gradual evaporation at a rate comparable to that of natural splash pools. After desiccation, we rehydrated both types of cells by adding artificial seawater and monitored their recovery. We found that desiccated sheets that had undergone rapid evaporation never gave rise to viable cells after rehydration (**Figure 4N**). In contrast, rehydrated cysts consistently gave rise to viable sheets (**Figure 4N**). These observations show that cysts, unlike flagellates, are equipped to survive hypersalinity and desiccation, and suggest that encystation confers a selective advantage during evaporation.

We further wondered whether multicellularity, by contrast, was advantageous in the other phase of the evaporation-refilling cycle, marked by low salinity. Multicellular choanoflagellates have been proposed to feed more efficiently, thanks to cooperative hydrodynamic interactions between flagella that enhance prey capture^60–62^, although attempts to test this hypothesis in *S. rosetta* have yielded conflicting results^60,63,64^. To test for a feeding advantage in multicellular *C. flexa* sheets, we quantified the capture of fluorescent bacteria in sheets and in dissociated single flagellates (**Figure 4O** and **S16B**). We found that sheets captured more than twice as many fluorescent bacteria per cell as single cells (**Figure 4O**). This suggests that multicellularity confers a prey capture advantage at salinities compatible with the maintenance of a functional collar complex, and therefore with feeding.

## Clonal-aggregative multicellularity is a versatile strategy for robust re-establishment of multicellularity in a variable environment

Although splash pools all undergo similar evaporation-refilling cycles, they vary in size, evaporation rate, and salinity after refilling. Given this variability, we wondered whether environmental parameters might shift the relative contributions of clonality and aggregation. Notably, we reasoned that conditions that impair cell division might favour aggregation, whereas conditions that limit cell-cell encounters might favour clonal multicellularity.

We first examined the influence of salinity within the range compatible with multicellularity (*i.e.,* below the dissociation-encystation threshold). At seawater salinity (1X), both cell division and aggregation occurred (**Figure 5A** and **S17A**). By contrast, medium-high salinity (2X) arrested cell division but did not affect aggregation (**Figure 5A** and **S17A**). Thus, *C. flexa* sheets formed through mixed clonal-aggregative multicellularity at 1X salinity, but through pure aggregation at 2X salinity.

**Figure 5.**
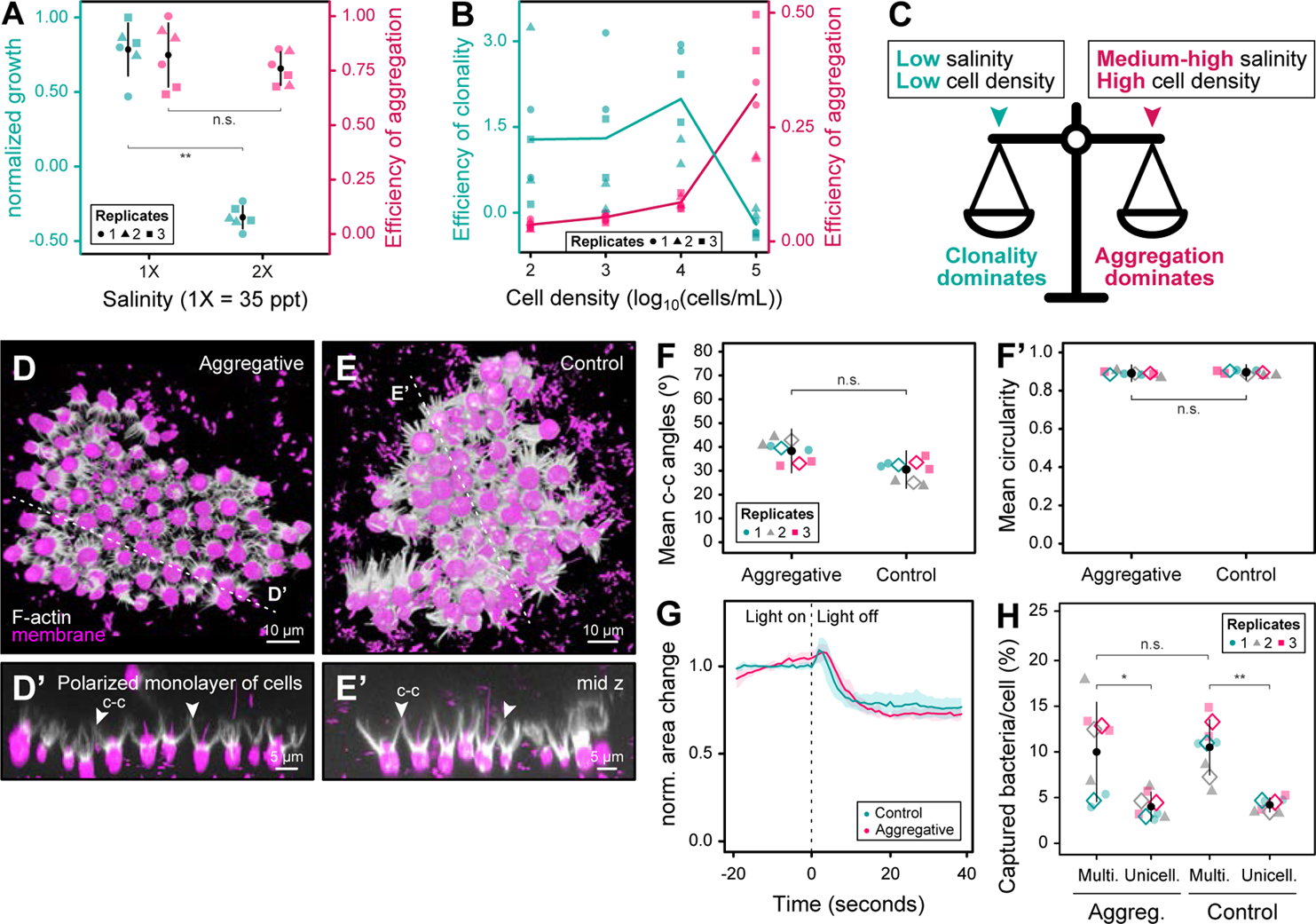
Environmental parameters modulate the relative contributions of clonality and aggregation. (A) Normalised growth of *C. flexa* cells at different salinities during gradual evaporation (turquoise, left y-axis; from experiment in Figure 4L). Efficiency of aggregation of *C. flexa* cells at different salinities (magenta, right y-axis), calculated based on the area of sheets in a 2-hour time course experiment (see Materials and Methods). Black circles: mean; error bars: standard deviation. (B) Efficiency of clonality (turquoise, left y-axis) and aggregation (magenta, right y-axis) at different initial cell densities, based on the area of sheets in a 24-hour time course experiment (see Materials and Methods). Lines: mean. (C) Schematics illustrating the relative contributions of clonality and aggregation to *C. flexa* multicellularity, as modulated by environmental conditions. At low salinity and low cell density, clonality dominates, whereas at medium-high salinity and high density, aggregation dominates. (D-E) 3D reconstruction of Airyscan confocal z-stacks of purely aggregative sheets (D) and control sheets (E), stained for membrane/cytoplasm (FM 4-64FX, magenta) and F-actin (phalloidin, white). (D’-E’) Mid z cross-sections (dashed lines) in D-E. Note that cells are connected by direct collar-collar contacts (c-c, white arrowheads), forming polarised monolayers of cells. (F) Quantification of mean collar-collar angles between cells in aggregative and control sheets in D-E. (F’) Quantification of mean circularity of aggregative and control sheets in D-E. Black circles: mean; error bars: standard deviation; diamonds: mean values of each independent replicate (n=41 aggregative sheets; n=46 control sheets). (G) Normalised colony area changes before and after light-to-dark transitions (vertical dashed line, t=0) in control or purely aggregative sheets. Line: normalised mean area; ribbon: standard deviation. n=5 sheets quantified by condition. (H) *C. flexa* aggregative and control multicellular sheets are more efficient at capturing bacteria than unicellular flagellates. Experiment performed as in Figure 4O. Black circles: means; error bars: standard deviations; diamonds: means of independent biological replicates. Statistics in A, F-F’, and H by the Mann-Whitney U test (* for p<0.05; ** for p<0.01; *** for p<0.001; n.s., non-significant). Figure related to **Figures S17-S19** and **movie S19**.

We next examined the effect of cell density, reasoning that it might influence aggregation efficiency by modulating encounter rates between cells. Field measurements across 12 splash pools revealed densities ranging from 30 to 4.4*10^4^ cells/mL (**Figure S17B; Table S1;** see Materials and Methods). This range overlapped with cell concentrations used in laboratory aggregation experiments (from 1.0*10^4^ to 6.7*10^5^ cells/mL), confirming that *C. flexa* can reach densities sufficient for aggregation in nature. To quantify the effect of cell density, we seeded dissociated cells at 10^2^ to 10^5^ cells/mL under both 1X and 2X salinity. We assessed aggregation efficiency by measuring final colony size under purely aggregative conditions (2X salinity), and clonality efficiency by quantifying additional colony growth at 1X salinity over the 2X baseline (see Materials and Methods). Measurements were taken during early sheet formation to avoid saturation effects. Aggregation efficiency increased monotonically with cell density and peaked at the highest density tested (10^5^ cells/mL) (**Figure 5B** and **S17C**). By contrast, clonality efficiency remained constant across intermediate densities (10^2^-10^4^ cells/mL) but decreased at the highest density (10^5^ cells/mL), likely due to depletion of bacterial prey limiting proliferation (**Figure 5B** and **S17C**). Therefore, low densities favour clonal multicellularity, high densities favour aggregative multicellularity, and intermediate densities support mixed clonal-aggregative formation.

Taken together, these data show that clonal-aggregative multicellularity spans a spectrum along which environmental conditions modulate the relative contributions of clonality and aggregation (**Figure 5C**). Importantly, certain environmentally relevant conditions – such as medium-high salinity or extreme cell densities – suppress one mode, resulting in purely clonal or purely aggregative multicellularity. This plasticity thus allows *C. flexa* to achieve multicellularity across a broader environmental range than either mechanism alone would permit.

## *C. flexa* sheets formed by aggregation are equivalent to control sheets in morphology and behaviour

Having identified environmentally relevant conditions that modulate the balance between clonal and aggregative multicellularity, we next asked whether *C. flexa* colonies formed by aggregation are functionally equivalent to those formed by cell division. We set out to characterise purely aggregative sheets, obtained by seeding dissociated cells at high density under 2X salinity. These were compared to control sheets generated under conditions permissive for clonality (1X salinity and a low initial density of 200 cells/mL; see Materials and Methods). Both control and aggregative sheets exhibited a similar polarised monolayer structure (**Figures 5D-E** and **S17D-G**) and were statistically indistinguishable in size (**Figure S17H**), proportion of aligned cells (**Figure S17I**), collar-collar angles (**Figure 5F** and **S17I’**), and circularity (**Figures 5F’** and **S17J**).

To test whether aggregation could also support the growth of pre-formed colonies independently of cell division, we fluorescently labelled single cells and added them to pre-formed sheets under 2X salinity (**Figure S18A**). The labelled cells initially adhered at the periphery of the sheets in variable orientations (**Figure S18B-C,F**) but, after 24 hours, they were robustly incorporated into the pre-existing colonies, displaying collar-collar contacts and aligned apico-basal polarity with their unlabelled neighbours (**Figure S18D-F**). These observations show that aggregation alone is sufficient to support both the formation and growth of sheets with wild-type morphology, independently of cell division.

We then characterised the behaviour of purely aggregative sheets. Both control and aggregative sheets exhibited equivalent inversion behaviour in response to light-to-dark transitions (**Figures 5G** and **S19**) and comparable prey capture efficiency over single cells (**Figure 5H**). In an independent assay, we also generated sheets by aggregation of cells labelled with two different colours (as in **Figure 2H**) and confirmed that the resulting chimeric sheets robustly inverted in response to light-to-dark transitions (**Movie S19**). These results demonstrate that *C. flexa* can form fully functional colonies through aggregation alone.

Thus, although environmental parameters modulate the strategy by which *C. flexa* becomes multicellular, the morphology and behaviour of the resulting sheets appear largely uncoupled from their formation mechanism.

## Aggregation is species-specific and constrained by kin recognition

To assess the relevance of aggregation in the life history of *C. flexa*, we tested for biological signatures of aggregative multicellularity. A frequent feature of this process is species specificity, whereby cells aggregate preferentially with their own species and little or not at all with others^65^. We tested this by mixing equal quantities of *C. flexa* and *S. rosetta* single cells. *C. flexa* formed aggregative sheets that completely excluded *S. rosetta*, demonstrating species-specific aggregation (**Figure S20**). Notably, *S. rosetta* cells did not aggregate with each other under these conditions, confirming that aggregation is specific to *C. flexa* rather than a general behaviour of choanoflagellate cells at sufficient density.

A second frequent feature of aggregative multicellularity is kin recognition – *i.e.,* the ability to discriminate kin versus non-kin and to preferentially aggregate with closely related strains within the same species. Kin recognition increases genetic homogeneity within the resulting multicellular entity, which is thought to facilitate the evolution of coordinated collective behaviour^66^ and/or restrict the spread of cheater mutants^16^. We assessed kin recognition among three *C. flexa* strains isolated from different splash pools: Strain 1 (originally isolated in 2019 and used in laboratory experiments above), and Strains 2 and 3 (both isolated in 2023 during Exped-B; **Figure 6A-B**). Each strain was established by manual isolation of a single sheet, followed by amplification in laboratory cultures (referred to as ‘single-sheet-bottlenecked’ cultures). Isolation of single cells from each single-sheet-bottlenecked culture allowed the establishment of clonal strains (referred to as ‘single-cell-bottlenecked cultures’ or ‘clones’).

**Figure 6.**
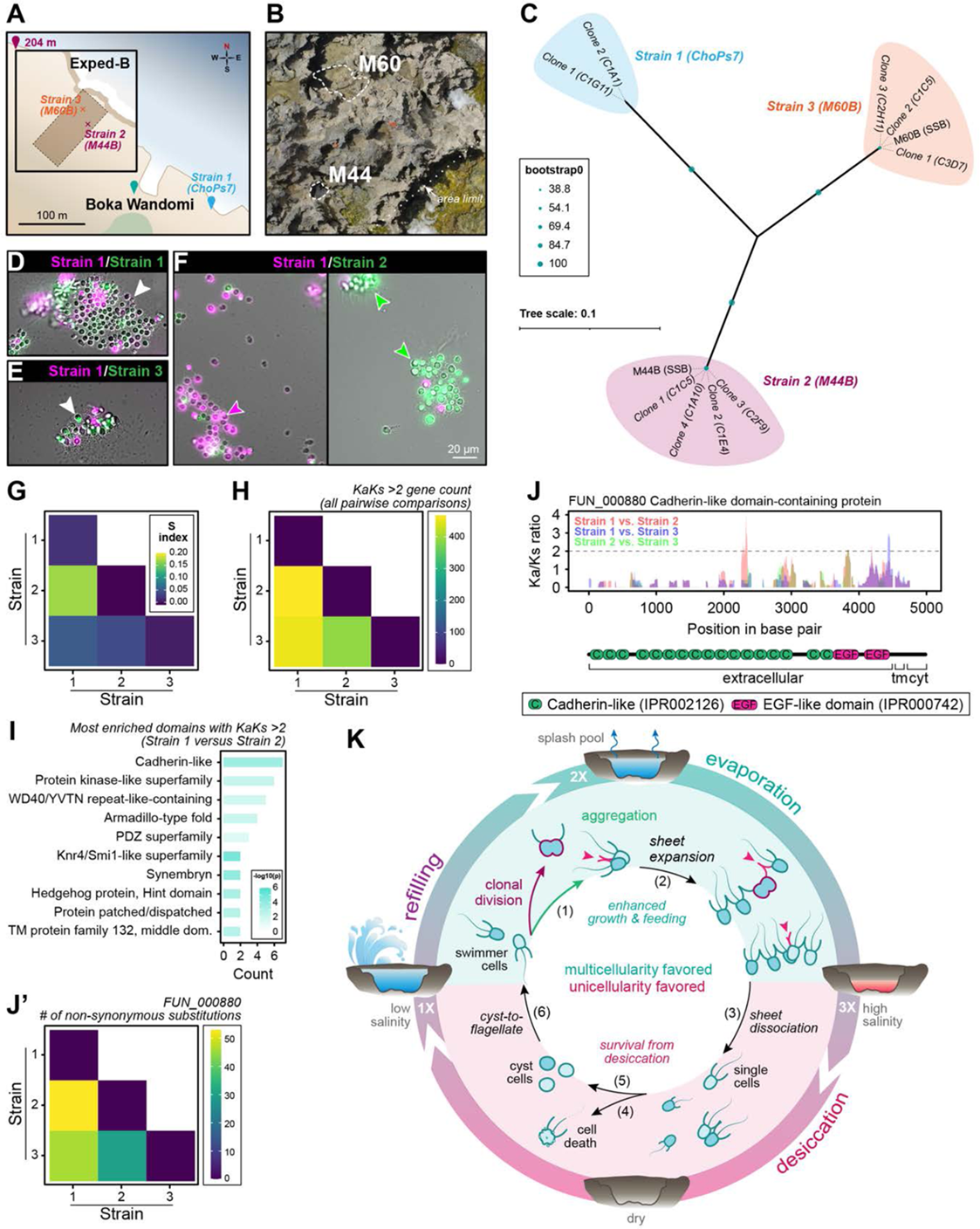
*C. flexa* aggregation is constrained by kin recognition. (A) Source location of the three distinct strains of *C. flexa* used in kin recognition experiments. (B) Aerial view of the M44 and M60 splash pools in the Exped-B site, from which Strains 2 and 3 were respectively isolated. Dashed lines: splash pool outlines. (C) Phylogenomic tree of Strains 1, 2 and 3, including single-sheet-bottleneck cultures (SSB) and three to four single-cell bottlenecked clones from each strain. (D) Cells of the same strain (here Strain 1), marked with magenta or green fluorophores, readily aggregated together regardless of colour, resulting in chimeric colonies (white arrowhead; compare Figure 2H). (E) Cells of Strains 1 and 3 (clone 1), marked with distinct fluorophores, readily aggregate into chimeric sheets (white arrowhead). (F) Cells of Strains 1 and 2 (clone 1), marked with distinct fluorophores, preferentially aggregate with their own strain, resulting in colonies with little or no chimerism (magenta and green arrowheads), indicating kin recognition. (G) Quantification of kin recognition using a segregation index (s) in different pairwise strain combinations (see **Figure S22** and Materials and Methods). The s index is calculated based on the proportion of green (or red) cells across sheets for a given combination of strains (as in^98^). An s index of 0 indicates no kin discrimination, while an s index of 1 indicates complete segregation of strains, and thus kin discrimination (**Figure S22B**, see Materials and Methods). *p* = 1.75e-09 by a one-way ANOVA. (H) Number of genes with high Ka/Ks regions (Ka/Ks > 2) for each pairwise comparison of strains. (I) Top 10 InterProScan domain annotations enriched in high Ka/Ks ratio regions in the Strain 1 versus Strain 2 comparison (p-value, Fisher’s exact test). Domain IDs: Cadherin-like (IPR002126); Protein kinase-like domain superfamily (IPR011009); WD40/YVTN repeat-like-containing domain superfamily (IPR015943); Armadillo-type fold (IPR016024); PDZ superfamily (IPR036034); Knr4/Smi1-like domain superfamily (IPR037883); Synembryn (IPR008376); Hedgehog protein, Hint domain (IPR001767); Protein patched/dispatched (IPR003392); Transmembrane protein family 132, middle domain (IPR031437). (J) Top candidate kin recognition gene encoding a cadherin-like domain-containing protein (FUN_000880) with a potential role in cell-cell adhesion. Upper row: Ka/Ks ratio along the gene coding sequence in each pairwise strain combination (red: Strain 1 vs. Strain 2; blue: Strain 1 vs. Strain 3; green: Strain 2 vs. Strain 3). Lower row: InterProScan domain architecture schematics. Tm: transmembrane domain; cyt: cytoplasmic domain. (J’) Number of non-synonymous substitutions in FUN_000880 for all pairwise strain combinations. (K) Summary schematic of *C. flexa* mixed clonal-aggregative multicellularity entrained by natural splash pool evaporation-refilling cycles. Multicellularity is favoured in low salinity, where sheets can form both by clonal division and by aggregation, and exhibit enhanced growth and prey capture (turquoise): (1) unicellular flagellate cells can divide clonally and/or aggregate to form multicellular sheets, maintaining direct cell-cell adhesions in their collar (magenta arrowheads); (2) sheets expand by clonal division, by aggregation of individual cells, and by sheet fusion. (3) Gradual evaporation leads to increase in salinity to medium-high levels, which prevents clonality but still allows aggregation and multicellularity. High salinity (>3-fold salinity) favours unicellularity (magenta): (4) *C. flexa* sheets dissociate into single cells that are incapable of proliferation and occasionally undergo cell death (5). Gradual evaporation, instead, results in the differentiation of flagellates into cysts (6) capable of surviving desiccation. (7) Rehydration restores permissive salinity and induces differentiation of cysts into flagellates, which can engage in clonal-aggregative multicellularity. Figure related to **Figures S20-S23, Table S3,** and **Supplementary Files S5-S6**.

To test for genetic divergence between these strains, we first sequenced, assembled, and annotated the *C. flexa* reference genome using a combination of long-read and Omni-C sequencing. The final assembly comprised 56 Mb and 14,084 genes across 528 scaffolds, with a BUSCO completeness score of 82.8%. This represents the third high-quality choanoflagellate genome after *M. brevicollis*^74^ and *S. rosetta*^67,68^. We then performed short-read sequencing of Strains 1, 2 and 3, as well as of three clonal descendants of each strain. Across all samples, we detected 193,564 SNPs, allowing us to reconstruct a phylogenomic tree in which all three parent strains were clearly delineated from each other (**Figure 6C** and **Supplementary Files S5,S6**). All clones clustered with their strain of origin with strong support and contained similar levels of genetic diversity to their corresponding single-sheet-bottlenecked strain (**Figure S21**). These results suggest either clonal formation of the originally isolated sheets (as in **Figure 1E-G**) or a loss of genetic diversity during sheet isolation and establishment of laboratory cultures (see Materials and Methods).

We tested kin recognition by dissociating sheets from all three strains, staining them with different fluorophores, and assessing aggregation across all pairwise strain combinations seeded at equal initial cell density (as in **Figure 2H**). All strains aggregated within hours, confirming that aggregation is a widespread behaviour among independently isolated *C. flexa* strains. Interestingly, cells of Strains 1 and 2 preferentially aggregated with their own strain when mixed, indicating kin recognition (**Figures 6D-G** and **S22**). By contrast, other pairwise combinations of strains showed no quantifiable preference for self-aggregation and readily formed chimeric sheets, as did cells of the same strain stained with different fluorophores (consistent with earlier results; **Figures 2H** and **S3**).

These results suggest the existence of a kin recognition system in *C. flexa*, reminiscent of those described in other aggregative multicellular organisms such as dictyostelid social amoebae. In the latter, kin recognition is mediated by polymorphic surface receptors rich in immunoglobulin and other adhesion domains^18,69^. In *Dictyostelium*, such receptors were first identified by their high ratio of non-synonymous to synonymous substitutions (Ka/Ks), a signature of diversifying selection^18^. We reasoned that, if *C. flexa* indeed possesses a comparable kin recognition mechanism, candidate kin recognition loci might be revealed by screening the genomes of Strains 1 and 2 for high Ka/Ks loci. Across all pairwise strain comparisons, we identified 840 genes with high Ka/Ks regions (**Figure 6H**), including multiple transmembrane proteins (**Figure S23A**). These high Ka/Ks genes encoded predicted proteins with a significant overrepresentation of domains consistent with adhesion (cadherins, TM132) or signalling (protein kinase) functions (**Figures 6I** and **S23B**). The most promising kin recognition candidates notably included one predicted cadherin (FUN_000880; **Figure 6I-J)** and one predicted receptor-protein kinase (FUN_003087; **Figures 6I** and **S23B-C**) that displayed particularly strong signatures of diversifying selection between Strains 1 and 2.

These findings indicate that polymorphic receptor candidates potentially mediating kin recognition can be readily identified in *C. flexa*, although some of these proteins may perform other functions, such as capture of bacterial prey. Determining their precise roles will require functional genetic tools, which are not yet available for *C. flexa*.

Taken together, species-specificity and kin recognition support the view that aggregation is a regulated biological process in *C. flexa*.

## Discussion

Our findings suggest that evaporation-refilling cycles of splash pools, the natural habitat of *C. flexa*, regulate its clonal-aggregative multicellularity (**Figure 6K**). *C. flexa* sheets are found in water-filled splash pools with salinities up to ∼3-fold that of natural seawater. As gradual evaporation proceeds, salinity increases, and sheets dissociate, with cells differentiating into solitary, non-motile, non-proliferative cysts. These cysts can survive complete desiccation and persist in the soil of the splash pools. Upon splash pool refilling, cysts regenerate a collar complex and transition back into free-swimming flagellates, which re-form multicellular sheets through aggregation and/or clonal division. This mixed mechanism confers two key advantages in this environment: (i) robust re-establishment of multicellularity across a broad range of conditions, and (ii) fast acquisition of multicellularity by simultaneous action of both mechanisms. Indeed, aggregation generally provides a speed advantage over clonality, as aggregative eukaryotes often complete multicellular formation within a few hours^20,20,70^, whereas clonal multicellularity is constrained by cell cycle duration, which ranges from one to eleven days in diverse micro-organisms in the wild (**Table S3**). Thus, the ability of *C. flexa* to aggregate may therefore allow faster and more robust re-establishment of multicellularity in the variable and ephemeral environment of splash pools.

The environmentally entrained life history of *C. flexa* parallels other recently described examples in which unicellular-to-multicellular transitions are modulated by fluctuating environmental parameters, such as salinity in cyanobacteria inhabiting brackish environments^71^ and periodic flooding in cave-dwelling bacteria^72^. A selective advantage for regulated life cycles in fluctuating environments is also supported by laboratory experiments in yeast^73^ and by theoretical models^5,74,75^. The environmentally entrained life cycle of *C. flexa* may thus allow it to combine both the benefits of multicellularity (enhanced filter-feeding, presumably supporting faster proliferation) with those of the unicellular cyst form (resistance to desiccation).

The ability of *C. flexa* to aggregate contrasts with prevailing views of choanoflagellates, which are canonically considered to exhibit strictly clonal multicellularity^25,39,76–78^ – like animals, their sister-group. Our study therefore reveals unexpected diversity in the modes of multicellularity within a pivotal clade for reconstituting animal origins. Notably, strictly clonal multicellularity has so far been experimentally demonstrated in only a single choanoflagellate species, the model *S. rosetta*^39^. This suggests that the diversity of choanoflagellate multicellularity warrants more systematic re-exploration. Although incomplete character-state mapping currently limits ancestral-state reconstruction, broader taxonomic sampling in the future will ultimately clarify whether clonal-aggregative multicellularity contributed to the origins of animals (as proposed by some authors^27^) or represents a specific innovation of the *C. flexa* lineage.

The mixed clonal-aggregative multicellularity of *C. flexa* was a surprise, given that clonal and aggregative multicellularity are often depicted as mutually exclusive in holozoans^2,79^ and eukaryotes in general^80–82^. However, although clonal-aggregative multicellularity has not previously been reported in close relatives of animals, nor in the context of a regulated life cycle to our knowledge, clonality and aggregation occasionally cooperate in the formation of other biological structures. These include bacterial biofilms^83^, experimentally evolved clusters of bacteria^84^, predator-induced groups of freshwater algae (which can even combine different species)^10^, clusters of budding yeast^85^, and certain syncytial amoebae^19^. While these diverse processes may not all represent *bona fide* multicellularity^86^ – since the resulting structures often lack multicellular-level adaptations such as controlled shape, size, or collective behaviour – they demonstrate that aggregation and incomplete cell division can coexist. Clonal-aggregative multicellularity may therefore be more widespread than currently appreciated.

Aggregation presents a well-known evolutionary challenge: a single aggregate can combine cells of different ancestries and potentially different genotypes, creating the potential for genetic conflict^87–89^ and, perhaps more importantly, limiting the coevolution of genes required for the emergence of complex multicellular behaviours^66,90^. The risks of chimerism can be mitigated if aggregation is restricted to close relatives – either actively, through kin recognition mechanisms, or passively, through a spatially structured environment limiting dispersal^91,92^. The latter might be relevant to *C. flexa*, as splash pools are collections of physically disconnected environments that might promote geographic divergence of genotypes, as shown for other organisms in similar environments (*e.g., Daphnia* metapopulations in tide pools^93^). Splash pools may thus have favoured the evolution of aggregative multicellularity by initially relaxing selection for strong kin recognition mechanisms (which *C. flexa* nonetheless appears to have eventually evolved). This scenario aligns with the emerging concept of ‘ecological scaffolding’, which proposes that patchy environments can facilitate the emergence of multicellularity by fostering local cooperation^94^. In the future, these questions will be informed by deeper characterization of dispersal, molecular kin recognition mechanisms, and natural genetic diversity at different scales (within and between colonies, within and between splash pools) in *C. flexa*.

The complex life history of *C. flexa* illustrates the phenotypic plasticity of close unicellular relatives of animals and thus lends additional support to the emerging concept of a pre-metazoan origin for complex life cycles^21,95^. Beyond this, our study establishes *C. flexa* as a powerful model to study the establishment of multicellularity in a close relative of animals within its natural context. This contrasts with other well-characterised facultatively multicellular holozoans, such as *S. rosetta*^22^ and *C. owczarzaki*^96^, which were each isolated only once from their natural environment over two decades ago, inevitably restricting studies of unicellular-to-multicellular transitions in these species to laboratory settings. In the future, we expect the continued dialogue between field and laboratory studies of *C. flexa* to continue clarifying key questions, such as the selective advantage(s) and ecological consequences of multicellularity versus unicellularity, the evolvability of clonal-aggregative behaviours under diverse selective pressures in diverse environments, and the molecular mechanisms of kin recognition.

## Data availability

The annotated reference *C. flexa* genome has been deposited on Zenodo under doi: 10.5281/zenodo.13837466. Raw short reads for Strains 1, 2 and 3 have been deposited on Zenodo under doi: 10.5281/zenodo.13837614.

## Code availability

The codes used for SNP analysis are deposited on Github: https://github.com/uhoro/RRRR_SNP_analysis

## Acknowledgements

We enormously thank Nicole King for support during this project and for providing useful feedback. We thank the personnel of the Shete Boka National Park in Curaçao for access to the field site. Special thanks to Tess Linden for help during preliminary field experiments and feedback on the manuscript, as well as to Geneviève Milon and Noah Whiteman for advice on laboratory and field experiments, respectively. Thanks to David S. Booth for providing JRR with access to microscopes and lab space during revision of this work after peer-review. Yashraj Chavhan, Seemay Chou, Stefania Kapsetaki, and Iñaki Ruiz-Trillo and members from the Multicellgenome lab that provided useful feedback on the preprint version of the manuscript. Marvin Albert, Stéphane Rigaud and Jean-Yves Tinevez at the Image Analysis Hub at the Institut Pasteur provided advice on image analysis. Pierre-Henri Commere and Sebastien Megharba from the Flow Cytometry platform at Institut Pasteur provided technical support with single-cell isolation of *C. flexa* by flow cytometry. We acknowledge the help of the HPC Core Facility of the Institut Pasteur for bioinformatic analyses. We thank Thomas Swale (Dovetail Genomics/Cantata) and Tara Rickman (Phase Genomics) for help with genome sequencing and assembly and for drafting the relevant Methods sections. We also thank members of Brunet and King laboratories for feedback and useful discussions and Jaime Ramirez and Kayla Dinshaw for technical support during preliminary experiments. NRR is supported by the European Union’s Horizon Europe research and innovation funding program under a Marie Skłodowska-Curie Actions grant (FlexAggon, grant agreement ID: 101106415). JRR is supported by a National Science Foundation Graduate Research Fellowship Program (grant no. 1752814) and the National Science Foundation Postdoctoral Research Fellowship in Biology (grant no. 2507925). UH is supported by the PPU program of the Institut Pasteur. Work in the King lab (TB, JRR, BTL, MCC) is supported by the Howard Hughes Medical Institute. JLS is a Howard Hughes Medical Institute Awardee of the Life Sciences Research Foundation. TB and work in the Brunet lab are supported by the Institut Pasteur (G5 package), the ERC (EvoMorphoCell, grant agreement ID: 101040745), the Vallee Foundation Inc, and the CNRS (UMR 3691). Views and opinions expressed are, however, those of the author(s) only and do not necessarily reflect those of the European Union or the European Research Council. Neither the European Union nor the granting authority can be held responsible for them.

## Author contributions

NRR, JRR, TB conceptualised the study and coordinated the work. MCC, BTL and TB performed preliminary fieldwork that identified the Shete Boka site that gave first insights in the *C. flexa* salinity range. NRR, JRR and TB collected fieldwork data included in this study. NRR, JRR and TB performed all laboratory experiments, strain isolations and bioinformatic analyses (except those specified below). BTL performed preliminary aggregation assays and aphidicolin treatments. YF optimised live imaging of dual-labelling aggregation experiments and performed growth curves under addition of salts under the supervision of NRR. JLS performed genome assembly and annotation. UH performed behavioural assays with technical support from NRR and SNP analyses with technical guidance from JLS. CC performed single-cell bottleneck isolations and generated the material for genome sequencing of Strains 1, 2 and 3. EATH and MAV provided support for fieldwork studies. EATH performed drone orthomap imagery. JRR wrote the first draft of the manuscript which was subsequently edited by NRR and TB. NRR made the figures with feedback from JRR and TB. TB supervised the study. All authors contributed to the review of the manuscript before submission for publication and approved the final version.

## Inclusion & ethics statement

All Curaçao researchers who supported or performed fieldwork are authors of the present manuscript.

## Materials and Methods

See detailed Materials and Methods in the **Supplementary Information** file.

## Supplementary Material

Supplementary **Figures S1-S23**, **Tables S2-S3** and legends for **Table S1**, **Supplementary Files S1-S6** and **Supplementary Movies S1-S19** are provided in the **Supplementary Information** file.

